# Metagenomic analysis reveals large potential for carbon, nitrogen and sulfur cycling in coastal methanic sediments of the Bothnian Sea

**DOI:** 10.1101/553131

**Authors:** Olivia Rasigraf, Niels A.G.M. van Helmond, Jeroen Frank, Wytze K. Lenstra, Matthias Egger, Caroline P. Slomp, Mike S.M. Jetten

**Affiliations:** Department of Microbiology, Radboud University Nijmegen, Nijmegen, The Netherlands; Netherlands Earth System Science Centre (NESSC), Utrecht, The Netherlands; Department of Earth Sciences, Utrecht University, The Netherlands; Soehngen Institute of Anaerobic Microbiology (SIAM), Radboud University Nijmegen, Nijmegen, The Netherlands

**Author notes:** Current address: German Research Centre for Geosciences (GFZ), Section 3.7 Geomicrobiology, Potsdam, Germany. Current address: The Ocean Cleanup, Rotterdam, The Netherlands (Matthias Egger).

## Abstract

The Bothnian Sea is an oligotrophic brackish basin characterized by low salinity and high concentrations of reactive iron, methane and ammonium in the sediments potentially enabling an intricate microbial network. Therefore, we analyzed and compared biogeochemical and microbial profiles at one offshore and two near coastal sites in the Bothnian Sea. 16S rRNA amplicon sequence analysis revealed stratification of both bacterial and archaeal taxa in accordance with the geochemical gradients of iron, sulfate and methane. The communities at the two near coastal sites were more similar to each other than that at the offshore site located at a greater water depth. To obtain insights into the metabolic networks within the iron-rich methanic sediment layer located below the sulfate-methane transition zone (SMTZ), we performed metagenomic sequencing of sediment-derived DNA. Genome bins retrieved from the most abundant bacterial and archaeal community members revealed a broad potential for respiratory sulfur metabolism via partially reduced sulfur species. Nitrogen cycling was dominated by reductive processes via a truncated denitrification pathway encoded exclusively by bacterial lineages. Gene-centric fermentative metabolism analysis indicated the central role of acetate, formate, alcohols and hydrogen in the analyzed anaerobic sediment. Methanogenic/-trophic pathways were dominated by *Methanosaetaceae*, *Methanosarcinaceae*, *Methanomassiliicoccaceae, Methanoregulaceae* and ANME-2 archaea. *Thorarchaeota* and *Bathyarchaeota* encoded pathways for acetogenesis. Our results indicate flexible metabolic capabilities of core community bacterial and archaeal taxa, which can adapt to changing redox conditions, and with a spatial distribution in Bothnian Sea sediments that is likely governed by the quality of available organic substrates.

## Introduction

Sediment microbial communities drive biogeochemical cycles through their specific metabolic activities. The supply of organic carbon from primary production or terrestrial input via rivers, and electron acceptors such as nitrate (NO_3_^−^) and sulfate (SO_4_^2−^) in marine systems, will select for particular microbial guilds. Together they will determine the establishment of environment-specific metabolic networks and geochemical profiles. Despite the critical role of coastal sediments in global biogeochemical cycling, for example, as a source of methane (CH_4_) (Bange et al. 1994) and sink for nutrients (Asmala et al. 2017), our understanding of their microbial community composition and how this is linked to the cycling of sulfur (S), carbon (C) and nitrogen (N), is still incomplete.

The Bothnian Sea, a brackish basin located in the northern part of the Baltic Sea, is an ideal location to study the linkage between microbes and biogeochemistry because of the distinct sharp redox zonation of its surface sediments (Egger et al. 2015a; Lenstra et al. 2018; Rasigraf et al. 2017). The Bothnian Sea is oligotrophic and most organic matter in the sediment is supplied through rivers and is thus of terrestrial origin (Algesten et al. 2006). SO_4_^2−^concentrations in the bottom water are low (3-5 mM), which has allowed the development of a relatively shallow SO_4_^2−^reduction zone in the sediment at sites with relatively high sedimentation rates. At such sites, CH_4_ is abundant in the lower part of the SO_4_^2−^ reduction zone, and a distinct sulfate-methane transition zone (SMTZ) has developed (Egger et al. 2015a; Lenstra et al. 2018). The exact position of the SMTZ varies with space and time depending on the sedimentation rate and the input of organic matter (Egger et al. 2015a; Lenstra et al. 2018; Rooze et al. 2016; Slomp et al. 2013). The input of reactive iron (oxyhydroxides, henceforth termed Fe oxides) is in general higher than sulfide (H_2_S) formation in the sediment resulting in net burial of Fe oxides below the SMTZ. Both modeling and incubation studies suggest CH_4_ oxidation with Fe oxides as the electron acceptor in the SO_4_^2−^-depleted methanic layers below the SMTZ (Egger et al. 2015b; Rooze et al. 2016; Slomp et al. 2013). So far, the underlying pathways and responsible organisms for this process are largely unknown.

Several studies have investigated the microbial community composition in sediments of the Bothnian Bay and North Sea with 16S rRNA pyrosequencing techniques and speculated on possible microbial guilds involved in CH_4_ and Fe cycling (Oni et al. 2015a; Reyes et al. 2016). In surface sediments from the Skagerrak and Bothnian Bay, various potential Fe-reducers belonging to *Desulfobulbaceae*, *Desulfuromonadaceae* and *Pelobacteraceae* families were identified (Reyes et al. 2016). In deeper methanic sediment layers of the Helgoland area in the North Sea, microbial populations predicted to be involved in Fe and CH_4_ cycling included uncultured lineages of candidate division JS1 and methanogenic/-trophic archaea belonging to *Methanohalobium*, *Methanosaeta* and anaerobic methane oxidizing archaea clade 3 (ANME-3) (Oni et al. 2015a). Moreover, recent findings indicate that temperature is another factor which can influence the pathway of crystalline Fe utilization in these sediments (Aromokeye et al. 2018). Investigations of microbial communities involved in Fe cycling are challenging due to the absence of suitable ‘universal’ biomarkers. Different microbial groups have evolved different mechanisms and underlying genes encoding responsible enzymes may be unrelated. Novel mechanisms with unknown enzymatic steps in Fe reduction may exist but would remain undetected.

Activity measurements and functional biomarker analysis showed the presence of various pathways for N cycling in the Bothnian Sea and Bothnian Bay sediments (Bonaglia et al. 2017; Hellemann et al. 2017; Rasigraf et al. 2017; Reyes et al. 2017). Thus, for example, dissimilatory nitrate reduction to ammonium (DNRA) and denitrification were shown to be of nearly equal importance in oligotrophic sediments at a coastal site in the Bothnian Bay (Bonaglia et al. 2017). These results contradict the common assumption that DNRA is of minor importance in oligotrophic sediments with low organic carbon input and low rates of H_2_S production. Also, a gene-centric approach for N-cycle potential was applied previously to the Bothnian Sea and Bothnian Bay sediments. Sediment from the surface layer, SMTZ and deep methanic zone were analyzed and showed that N cycling genes were most abundant in the surface layer with denitrification being potentially the dominant pathway for N loss (Rasigraf et al. 2017). Furthermore, in suboxic sediments from the Bothnian Bay, the genetic potential for denitrification was far greater than that for DNRA (Reyes et al. 2017). With respect to the nitrification potential, differences were found between analyzed sites in the Bothnian Sea and Bothnian Bay. While in the central part of the Bothnian Sea, the nitrification potential was almost exclusively attributed to ammonia oxidizing archaea (AOA) belonging to *Thaumarchaeota* Marine Group-I (MG-I) (Rasigraf et al. 2017), in suboxic coastal sediments in the Bothnian Bay, both AOA and ammonia oxidizing bacteria (AOB) seemed equally important (Reyes et al. 2017). Thus, large differences in measured activities and genetic inventory can occur between sediments in the same region. The environmental factors driving those differences are not well explored.

Here, we assessed the microbial community composition in sediments at three sites along a water depth gradient in the Bothnian Sea by various complementary approaches including 16S rRNA amplicon sequencing. Two sites are located near the coast in the Öre Estuary (N10 and NB8, Lenstra et al. 2018) while the third site is located in the central-basin of the Bothnian Sea (US5B, Egger et al. 2015a). Porewater profiles of key geochemical constituents such as SO_4_^2−^, dissolved Fe and CH_4_ were used to determine the redox zonation. In addition to community comparisons between sites, we examined the core microbial community and its metabolic potential in the ferruginous methanic zone at one of the coastal sites through metagenome sequencing. Several high quality metagenome-assembled genomes (MAGs) were recovered for abundant microbial community members. Analysis of the MAGs indicated a flexible metabolic network with a strong potential for fermentation and S cycling.

## Materials and Methods

### Sampling and geochemical analysis

The two near-coastal sites, N10 and NB8 are located in the Öre Estuary in the Bothnian Sea at water depths of 21 and 33 m, respectively (Lenstra et al. 2018). Sediments at these sites were collected during a field campaign with R/V *Lotty* in August 2015 using a Gemini gravity corer (8 cm inner diameter). The offshore site US5B is located in the central basin of the Bothnian Sea at a water depth of 214 m and was sampled in August 2012 as described in Egger et al. 2015a. Sediments at this site were collected during a field campaign with R/V *Aranda* in August 2012 using a GEMAX gravity corer (10 cm inner diameter). Locations of all sampled sites are shown in Figure 1. Porewater depth profiles of SO_4_^2−^, CH_4_, NH_4_^+^, H_2_S and dissolved Fe were measured either onboard or later in the laboratory as described previously (Egger et al. 2015a; Lenstra et al. 2018; Figure 2). Sediment characteristics of sampled sites are summarized in Table 1. Sediment cores were kept at 4°C in the dark covered with a water layer until slicing. The slicing of sediment cores was performed in an anaerobic chamber under argon atmosphere. Sediment subsamples dedicated for DNA isolation were stored at −20°C until further processing.

**Table 1:** Characteristics of the investigated sites N10, NB8 and US5B in the Bothnian Sea. The data for water depth, temperature, coordinates, organic carbon content and sedimentation rates were compiled with from Lenstra et al. 2018 and Egger et al. 2015a. Abbreviations: mbss, meters below sea surface.

| Site | Water depth<br>(mbss) | Temperature<br>°C | Latitude<br>°N | Longitude<br>°E | C<sub>org</sub> (wt. %) | Sed. rate (cm<br>year<sup>-1</sup>) |
| --- | --- | --- | --- | --- | --- | --- |
| N10 | 20.8 | 7.8 | 63.293 | 19.462 | 3.40 (± 0.70) | 0.25 |
| NB8 | 33.2 | 6.3 | 63.291 | 19.495 | 3.85 (± 0.07) | 1 |
| US5B | 214 | 5.0 | 62.351 | 19.581 | 2.58 (± 0.21) | 1.3 |

**Figure 1:**
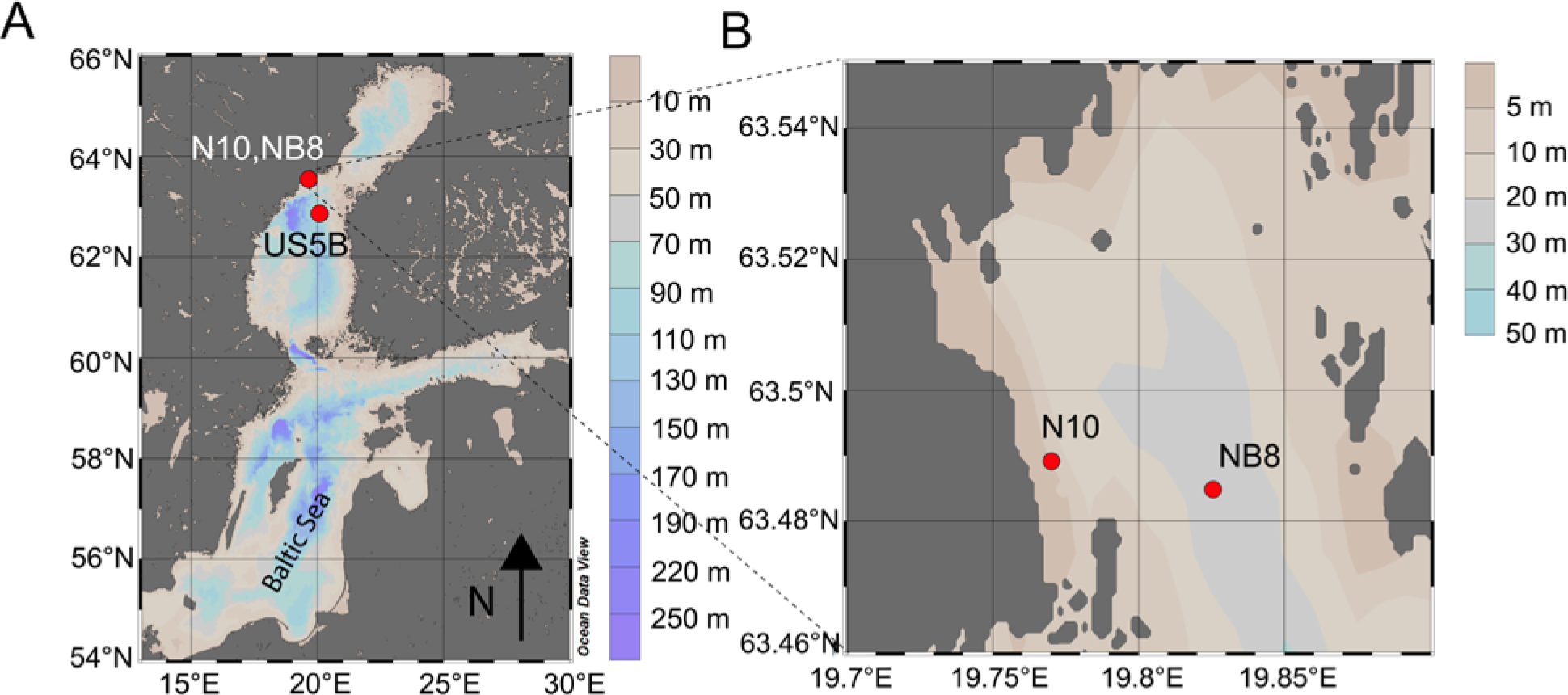
(a) Locations of sampling sites N10, NB8 and US5B in the Bothnian Sea; (b) Locations of both coastal sites N10 and NB8 in the Öre Estuary in the Bothnian Sea. Figure drawn using Ocean Data View (Schlitzer 2015).

**Figure 2:**
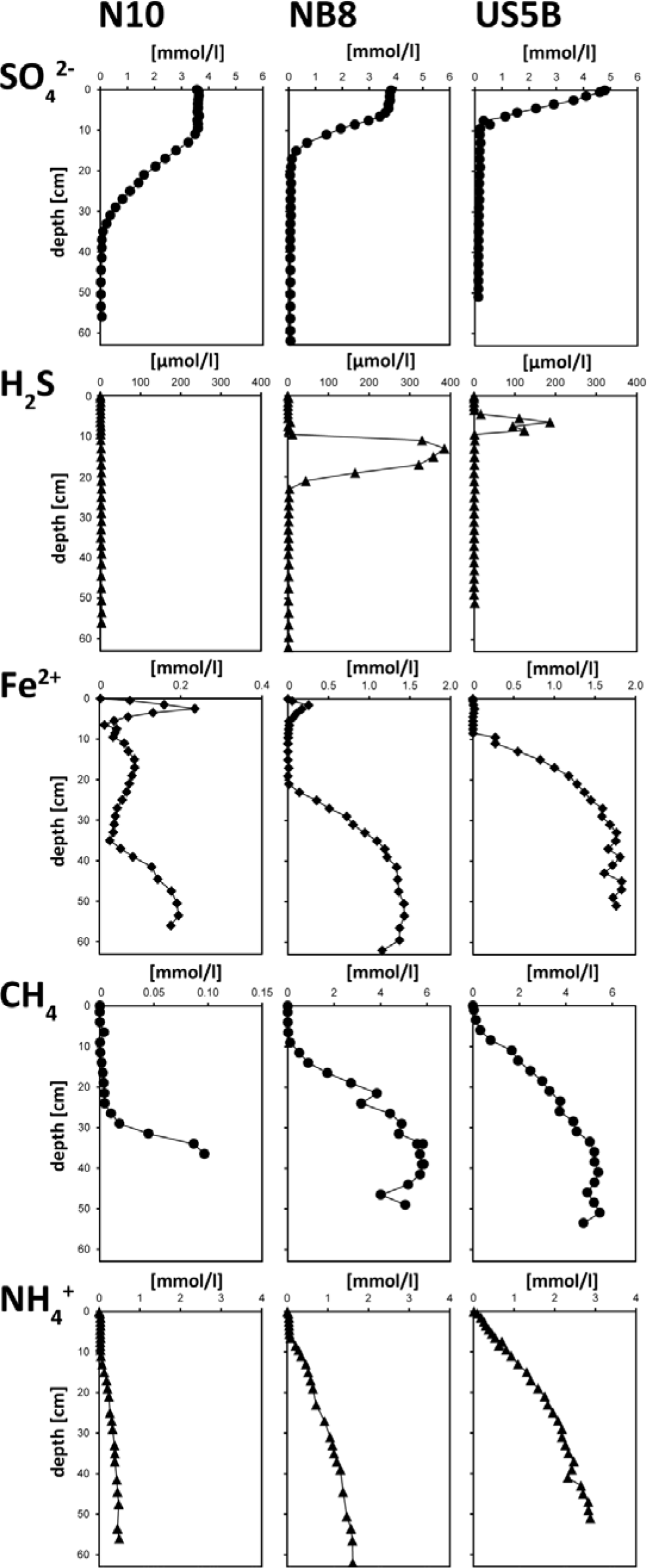
Geochemical profiles for sites N10, NB8 and US5B. Porewater profiles of SO_4_^2−^, sulfide (∑H_2_S=H_2_S + HS^−^ + S^2−^), Fe^2+^, CH_4_ and NH_4_^+^ are shown. Note the different scales for site N10.

### DNA isolation

The frozen core sediment subsamples were defrosted on ice and vortexed to obtain homogenous slurry. Subsequently, 0.2-0.5ml of original sediment slurry was filled into a bead beating tube from the PowerSoil DNA isolation kit (MoBio, USA). Further isolation was performed according to manufacturer’s instructions. The quantity of isolated DNA was assessed by NanoDrop 1000 (Thermo Scientific, USA) and Qubit^®^ 2.0 (Invitrogen, Life Technologies, Carlsbad, USA). After isolation, DNA was frozen at −20°C until further use.

### 16S rRNA and metagenome sequencing

The amplification of total archaeal and bacterial 16S rRNA genes was performed with the following primer pairs: Arch349F (5’-GYGCASCAGKCGMGAAW30) (Takai and Horikoshi 2000) and Arch806R (5’-GGACTACVSGGGTATCTAAT-3’) (Takai and Horikoshi 2000) for archaea, Bac341F (5’-CCTACGGGNGGCWGCAG-3’) (Herlemann et al. 2011) and Bac806R (5’-GGACTACHVGGGTWTCTAAT-3’) (Caporaso et al. 2012) for bacteria. 16S rRNA amplicon sequencing was performed on the Illumna MiSeq platform using the MiSeq Reagent Kit v3, yielding 2x300bp paired-end reads (Macrogen Inc., Europe).

For metagenomic sequencing, DNA from separate depth samples was pooled in equimolar concentrations. Paired-end metagenomic sequencing with 2x300bp sequence chemistry was performed with Miseq reagent kit v3 on Illumina Miseq platform (San Diego, California, USA) according to manufacturer’s instructions at the Microbiology Department of Radboud University, Nijmegen.

### 16S rRNA gene amplicon analysis

Paired end reads were processed with the Mothur v.1.36.1 software following the standard operation procedure (MySeq SOP) instructions (Kozich et al. 2013). The length of overlapped sequences was filtered for 400-500 base pairs (bp). Chimeric sequences were removed with the UCHIME algorithm (Edgar et al. 2011). Sequences were clustered into operational taxonomic units (OTU) with a 97% identity cut‐off and classified using the SILVA 16S rRNA gene non‐redundant reference database (version 123, SSURef123NR99) and the Bayesian classifier (‘wang’) (Pruesse et al. 2007). After quality trimming, chimera removal and normalization (“subsampling” in Mothur) of data, each sample contained 5,000 sequences for bacteria and 2,000 sequences for archaea. Samples with fewer sequences were excluded from the analysis.

Statistical analysis was performed in R (https://www.r-project.org/) (R Development Core Team, 2013) with OTU tables obtained in Mothur using the package Vegan (Oksanen et al. 2018). Data visualization was performed in Rstudio (RStudio Team 2015) using the package ggplot2 (Wickham 2016). The R package “OTUtable” was used to merge identical taxonomic groups classified as different OTUs in Mothur (Linz et al. 2017).

### Metagenome analysis: assembly, binning, annotation

Sequencing data obtained from the sediment sample described in this study were analyzed together with data obtained from incubation samples which are part of another study (data not shown).

Quality-trimming, sequencing adapter removal and contaminant filtering of Illumina paired-end sequencing reads was performed using BBDuk (BBTools suite version 37.17) (Bushnell), yielding 97,703,456 reads. Processed reads were co-assembled using MEGAHIT v1.1.1-2 (Li et al. 2015; Li et al. 2016) using the “meta-sensitive” preset. MEGAHIT iteratively assembled the metagenome using k-mers of length 21, 29, 39, 59, 79, 99, 119, 141. Reads were mapped back to the assembled metagenome for each sample separately using Burrows-Wheeler Aligner 0.7.15 (Li and Durbin 2010) (BWA), employing the “mem” algorithm. The sequence mapping files were processed using SAMtools 1.6 (Li et al. 2009). Metagenome binning was performed for contigs greater than 2,000 bp. To optimize binning results, four different binning algorithms were used: COCACOLA (Lu et al. 2017), CONCOCT (Alneberg et al. 2014), MaxBin 2.0 2.2.3 (Wu et al. 2016) and MetaBAT 2 2.10.2 (Kang et al. 2015). The four bin sets were supplied to DAS Tool 1.0 (Sieber et al. 2018) for consensus binning to obtain the final bins. The quality of the genome bins was assessed through a single-copy marker gene analysis using CheckM 1.0.7 (Parks et al. 2015). A coarse taxonomic classification of the genome bins was performed using CheckM and further refined by placing bins in a phylogenetic tree using the UBCG pipeline for phylogenomic tree reconstruction (Na et al. 2018). Annotation and biomarker detection was performed with KEGG automatic annotation server with bit score threshold of 100 (Moriya et al. 2007) and the Microbial Annotation and Analysis Platform of MicroScope (MAGE) (Vallenet et al. 2006). All sequencing data obtained for this project were submitted to the GenBank under the BioProject PRJNA511814. The metagenome originating from the in situ sediment in the methanic Fe-rich zone at site NB8 described in this study is designated as sample BS5 (BioSample SAMN10644131).

### Metagenome analysis: *mcrA* biomarker analysis

Functional biomarker analysis was performed as described previously (Lüke et al. 2016; Rasigraf et al. 2017). Following the procedures described in Lüke et al. 2017, metagenome data for the in situ sediment sample were quality trimmed with CLC Genomics Workbench 9.5.3 software using the following settings: quality score limit 0.01 (Q20), maximum number of ambiguous base pairs 0, min read length 100 [nt]. Metagenome size comprised 18,107,912 reads after quality trimming. Functional biomarkers were identified with blastx (release 2.4.0) using manually curated functional gene databases following the procedure described previously. Amino acid sequence data were aligned in ARB (Ludwig et al. 2004) and used for building an alternative classification taxonomy in MEGAN 5.11.3 based on manually curated *mcrA* gene database (Huson et al. 2007). Curated functional gene reads were re-blasted with a database file adapted for alternative taxonomic classification in MEGAN. Blast output was then imported into MEGAN and visualized for quantitative analysis. In total 288 *mcrA* gene reads were extracted from the metagenome. Quantified data were visualized with the R statistical package ggplot2.

For quantitative comparison, the analyzed gene reads were normalized to metagenome size and average gene length according to the following formula: normalized read count = (gene read count*1,000,000,000)/(total metagenome read count*average gene length [nt]).

For SSU rRNA quantification, raw reads were mapped to SILVA database (release 128) in CLC Genomics Workbench with the following settings: match score 1, mismatch cost 2, insertion cost 3, deletion cost 3, length fraction 0.5, similarity fraction 0.8. Mapped reads were extracted and submitted to SILVAngs online analysis pipeline (www.arb-silva.de/ngs/).

## Results and Discussion

### Geochemistry of sites N10, NB8 and US5B

Porewater profiles revealed the presence of a shallow SMTZ at all three sites (Figure 2). At site N10, the SMTZ is located at a depth of about 25-35 cm. At this site, CH_4_ and H_2_S concentrations in the porewater were very low. At sites NB8 and US5B, in contrast, the SMTZ was located at depths of about 20-25 cm and 4-9 cm, respectively, and distinct maxima in H_2_S was observed within the SMTZ. Maximum concentrations of NH_4_^+^ at depth in the sediment ranged from about 0.5 mM at site N10 to 1.5 and 3.0 mM at sites NB8 and US5B, respectively. All sediments were rich in dissolved Fe^2+^, with concentrations increasing in the sequence N10, NB8 and US5B below the SMTZ (Figure 2). This spatial trend was in accordance with the observed 10-fold increase in sediment accumulation rates and corresponding increased input of organic matter with distance from the coast.

### Sediment microbial diversity in the Bothnian Sea

Bothnian Sea vertical sediment profiles were analyzed for their bacterial and archaeal populations at sites N10, NB8 and US5B. Results are presented in Figures 3 and 4.

**Figure 3:**
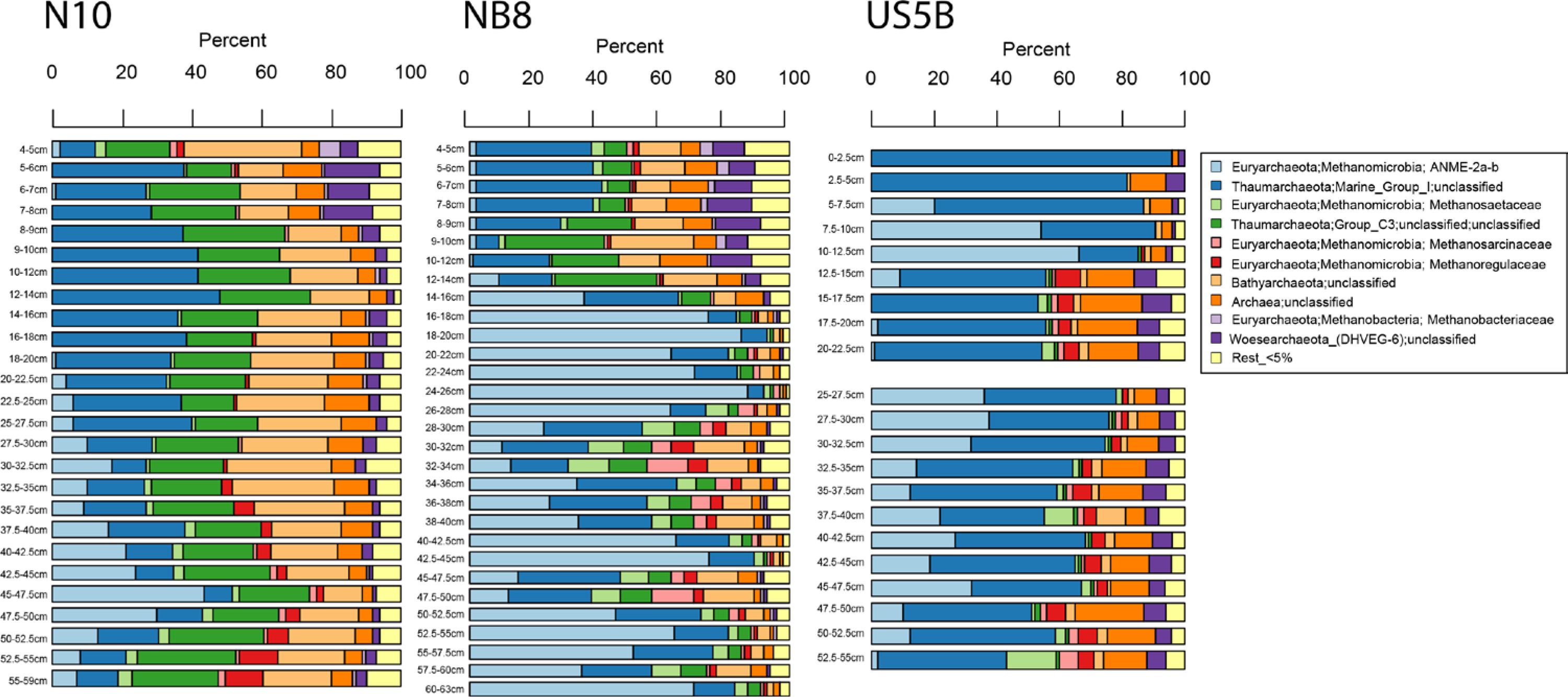
Distribution of archaeal taxons based on 16S rRNA amplicon analysis for sediment transects from N10, NB8 and US5B sites in the Bothnian Sea.

**Figure 4:**
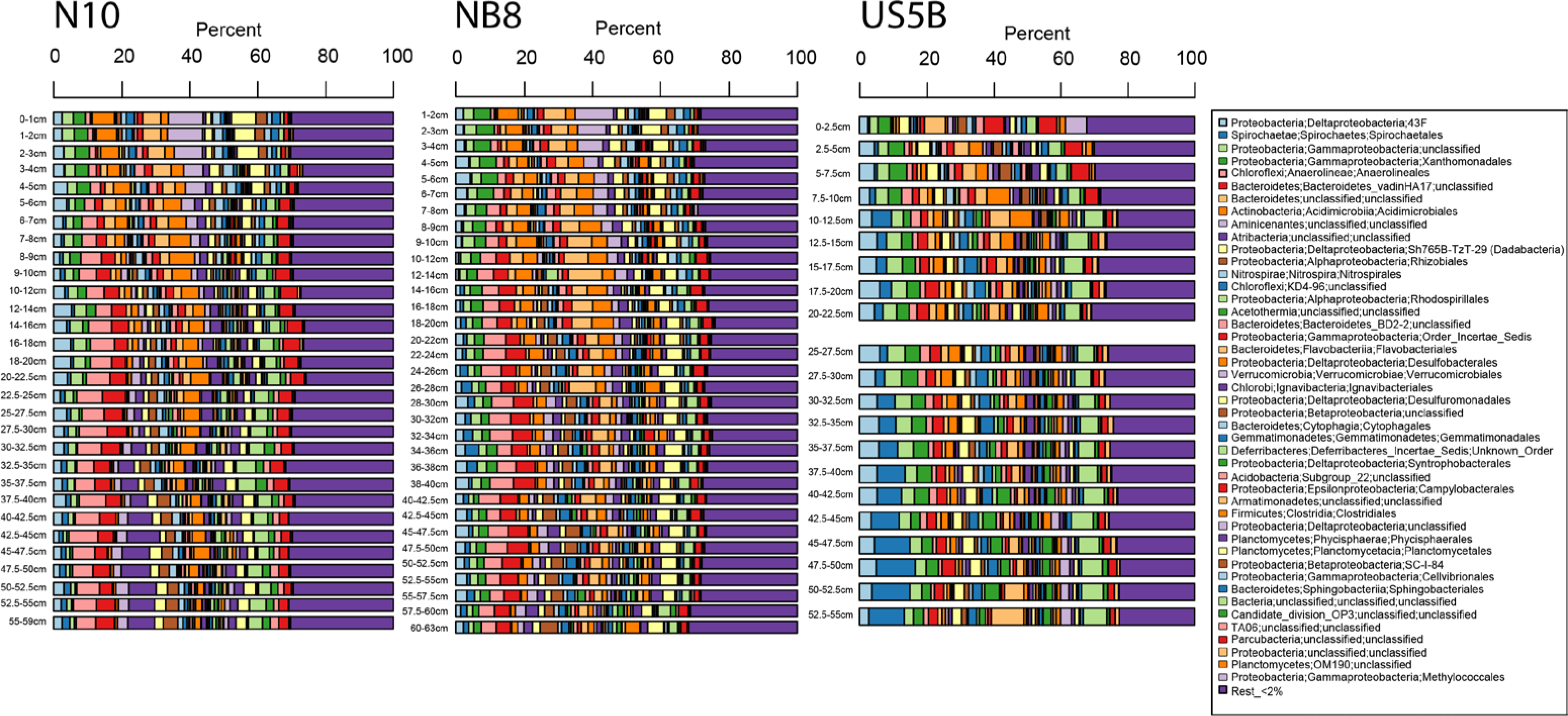
Distribution of bacterial taxons based on 16S rRNA amplicon analysis for sediment transects from N10, NB8 and US5B sites in the Bothnian Sea.

### Archaeal 16S rRNA gene distribution in the Bothnian Sea sediments

The vertical distribution of archaeal 16S rRNA gene sequences at all three sites over the sediment profile is shown in Figure 3. The results revealed that archaeal communities were more similar between the neighboring coastal sites N10 and NB8, than to the offshore site US5B. Despite the very similar geochemical profiles at sites NB8 and US5B, the communities were significantly different. Non-metric multidimensional scaling (nMDS) analysis performed on the archaeal species abundances from all sites revealed a clear separation as seen in Figure 5 (B).

**Figure 5:**
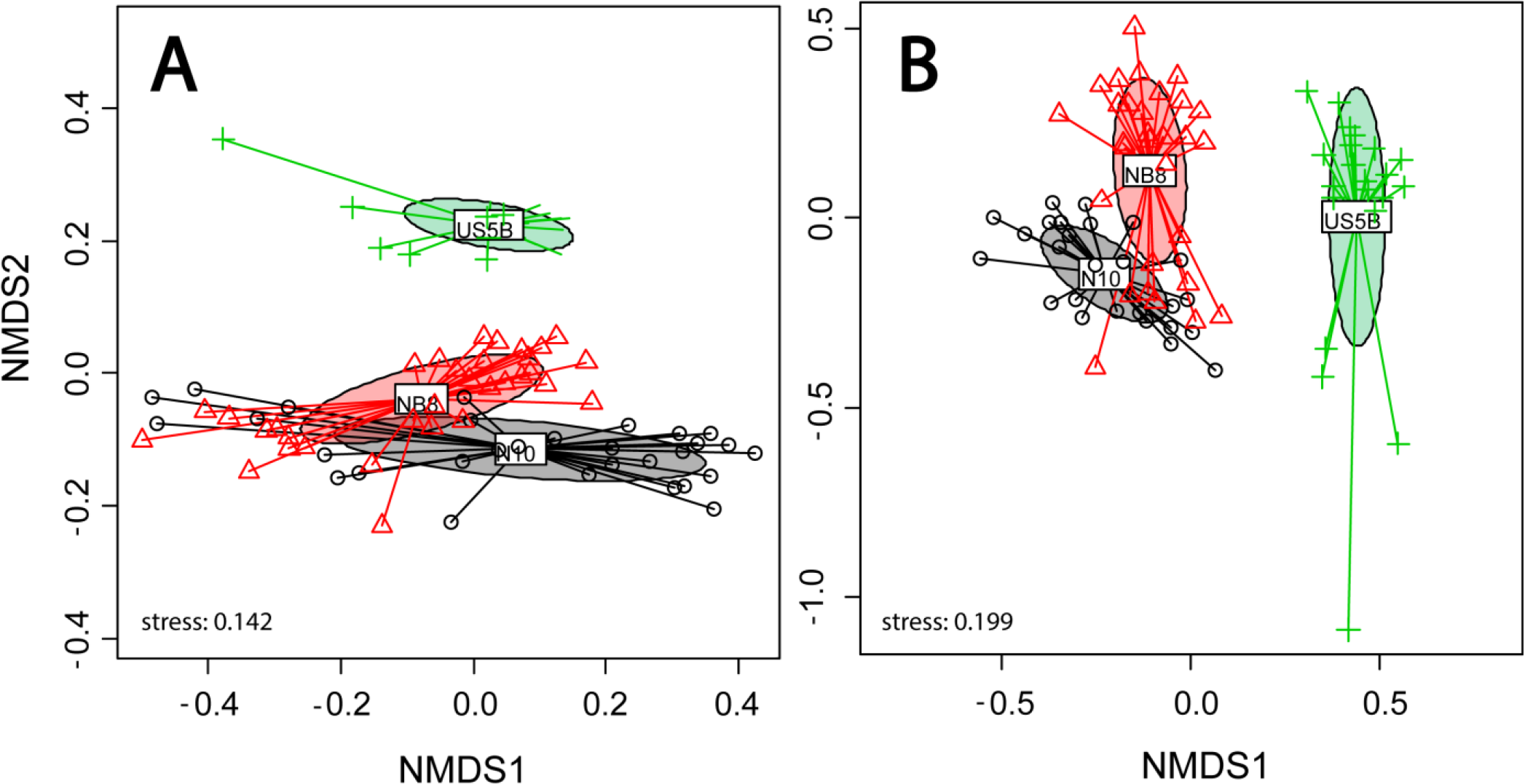
Non-metric multidimensional scaling (nMDS) analysis of sedimentary community populations from sampling sites N10, NB8 and US5B of (a) bacteria and (b) archaea based on 16S rRNA gene amplicon data. A dissimilarity matrix was calculated using the Bray-Curtis dissimilarity index in R. Symbols represent individual sampling depths of each sampling site, colors different sampling sites.

At all sites, upper sediment microbial communities were distinct from the ones at greater depth. This shift in relative species abundances could be explained by the rapid decrease in available electron acceptors, in particular SO_4_^2−^, and the increase of dissolved Fe and CH_4_ with depth. When comparing the archaeal community at different depths in the sediment, a similarity was observed for the upper sediment layer (4-18 cm) of site NB8 with the total profile of site N10. Since the upper sediment layer of site NB8 was characterized by a similar geochemistry as the whole profile of site N10, the observed similarity might indicate that the local microbial community was adapted to the prevailing environmental conditions.

At all sites, the upper SO_4_^2−^ containing sediment zone was dominated by *Thaumarchaeota* Marine Group-I (MG-I). Although their relative abundance decreased with depth, they still could be detected throughout the whole sediment profile at each site. MG-I have previously been shown to be ubiquitous in terrestrial and marine environments in which they are assumed to be involved in aerobic ammonia oxidation (Pester et al. 2011). However, several recent studies have detected genetic signatures of MG-I in deeper anaerobic sediment layers where an aerobic lifestyle is not likely (Rasigraf et al. 2017; and references therein). Some members of MG-I have been shown to use organic N compounds for growth, without possessing the aerobic ammonia oxidizing enzyme complex and the ability for ammonia oxidation (Weber et al. 2015).

Methanotrophic archaea assigned to the ANME-2a/b clade were the most dominant community member of *Euryarchaeota* and were detected at all analyzed sites. No other ANME clades could be detected. Their relative abundance peaked at the putative SMTZ zone (characterized by measurable porewater sulfide content). Below the SMTZ, ANME 16S rRNA biomarker showed a more scattered distribution. The ANME-2a/b archaeal clade has been detected in many marine and brackish sediments, including different parts of the Baltic Sea (Treude et al. 2005). However, some studies have shown its preference for shallow sediment depths with low CH_4_ and H_2_S concentrations (Roalkvam et al. 2012; Roalkvam et al. 2011). Such conditions are also found in the Bothnian Sea, where the salinity of the overlying water and H_2_S concentrations in the sediments are relatively low. Despite their central role in sulfur cycling, ANME-2a have also been linked to the oxidation of CH_4_ in the absence of SO_4_^2−^ by a direct electron transfer onto artificial shuttles (McGlynn et al. 2015; Scheller et al. 2016). Whether the Bothnian Sea ANME-2 organisms are able to use Fe oxides as an electron acceptor has not been shown so far. Previous research has indicated that Fe oxides stimulate CH_4_ oxidation in methanic sediments at site US5B (Egger et al. 2015b).

Other abundant archaeal groups comprised *Bathyarchaeota*, *Woesearchaeota* and *Thaumarchaeota* Group C3 (G-C3). Both, *Bathyarchaeota* and *Thaumarchaeota* G-C3 were relatively less abundant at US5B ranging between 1-9% and 1-2% of total archaeal 16S rRNA gene reads, respectively. Both groups were most prevalent in sediment layers above the SMTZ at NB8 (11-26% of all archaeal 16S rRNA reads for *Bathyarchaeota* and 7-32% for *Thaumarchaeota* G-C3) and throughout the whole profile of N10 (11-33% for *Bathyarchaeota* and 13-29% for *Thaumarchaeota* G-C3). Differences in sedimentation rates and quality of deposited organic matter which would differ between coastal and offshore sites, would both affect the degradation dynamics and intermediary metabolites. Thus, the prevalence of both groups could reflect the quality of degradable/fermentable organic matter in the same ecosystem such as the Bothnian Sea. For *Bathyarchaeota* several MAGs have been analyzed recently. Some studies found genomic indications for possible methylotrophic methanogenesis or anaerobic methanotrophy (Evans et al. 2015; Harris et al. 2018), others speculated on involvement in detrital protein degradation, fermentative acetate production and no capacity for methanogenesis (He et al. 2016; Lazar et al. 2016; Lloyd et al. 2013). *Thaumarchaeota* G-C3 16S rRNA gene sequences have been detected previously in a variety of terrestrial and marine environments (Hugoni et al. 2015; Na et al. 2015; Zeng et al. 2017). Their role remains somehow enigmatic since no enrichments or genomic sequences are available yet. Some members have been shown, however, to be involved in acetate consumption in SO_4_^2−^-reducing marine and estuarine sediments (Na et al. 2015; Webster et al. 2010).

At all sites, a significant fraction of archaeal reads could not be classified based on the database we used (SILVA release 123). For the deeper layers of US5B, this fraction ranged from 2 to 22% of the total archaeal 16S rRNA gene reads. However, the analysis of MAGs obtained from the methanic zone from site NB8 indicated that at least part of those unclassified archaea belonged to the newly described phyla *Thorarchaeota* and *Lokiarchaeota*. Based on the previously published results, *Thorarchaeota* have been discussed to be involved in acetate production and sulfur cycling by thiosulfate/elemental sulfur reduction (Seitz et al. 2016).

The distribution of *Woesearchaeota* at NB8 followed that of *Bathyarchaeota* and *Thaumarchaeota* G-C3. The highest relative abundance was observed above the SMTZ with 5-14% of total archaeal 16S rRNA gene reads. The abundance decreased below 1% max below the SMTZ. At US5B, the distribution of *Woesearchaeota* 16S rRNA gene reads was more even over the sediment depth profile with abundances ranging between 1-9% of total reads. At N10, *Woesearchaeota* were also more prevalent in the upper sediment layers above 8 cm with abundances reaching up to 16% of total reads, however they were also present throughout the whole core with 1-5% of total reads. The phylum *Woesearchaeota* was proposed in 2015 when first bins from environmental metagenomes were analyzed (Castelle et al. 2015). Small genomes and incomplete gene sets necessary for glycolysis, pentose phosphate pathway and pyruvate metabolism were discussed to be indicators for a symbiotic or parasitic lifestyle (Castelle et al. 2015). No woesearchaeal metagenomic bins could be retrieved from the analyzed depth at site NB8. This could be explained by a relatively low abundance of 16S RNA genes of *Woesearchaeota* in deeper layers of NB8 pointing to their low importance in those sediments.

Most abundant known methanogens from the phylum *Euryarchaeota* were represented by families *Methanosaetaceae*, *Methanosarcinaceae*, *Methanoregulaceae* and *Methanobacteriaceae*. The lowest proportional abundance of all methanogens was detected in SMTZ sediment layers. *Methanosaetaceae* were more prevalent in deeper layers of all sediment profiles with lowest numbers detected at N10. Their relative abundance reached 4% of total archaeal reads at N10, 16% at US5B and 13% at NB8. *Methanosarcinaceae* were present at all sites, with lowest numbers at N10 (2% max of total archaeal reads). Their distribution below the SMTZ at NB8 was rather scattered ranging between 1-13% of total archaeal reads. Higher abundances of *Methanosarcinaceae* correlated with lower abundances of ANME reads. At US5B, the abundance of *Methanosarcinaceae* reads ranged between 1-6% and was the highest in the deepest analyzed depth at 52.5-55 cm. This depth was also characterized by the highest observed proportion of *Methanosaetaceae* gene reads. *Methanoregulaceae* were more prevalent in sediment layers below the SMTZ at both US5B and NB8. At N10, their distribution correlated well with that of *Methanosaetaceae*. Relative abundances of total archaeal reads reached 8% at US5B, 7% at NB8 and 11% at N10. The distribution of *Methanobacteriaceae* was opposite to that of other methanogens. Highest abundances were detected above the SMTZ at NB8 with 4% max of total archaeal reads. Below the SMTZ, the abundance never exceeded 1% of total archaeal reads. At US5B, *Methanobacteriaceae* reads were below 1% throughout the sediment core. Also at N10, the highest abundance with 6% of total archaeal reads was observed in the upper most layer of 4-5 cm. In the deeper profile of N10, their abundance did not exceed 1% of total reads. Previous research has shown that *Methanosaetaceae* methanogens are strict acetotrophs and adapted to low acetate concentrations (Jetten et al. 1992). Low concentrations would be indicative of either low production or high turnover of acetate in Bothnian Sea sediments. In contrast, *Methanosarcinaceae* methanogens are generalists by being able to utilize a variety of substrates for CH_4_ production, but appear to possess lower affinities to acetate (Jetten et al. 1992; Liu and Whitman 2008). Both *Methanobacteriaceae* and *Methanoregulaceae* have been shown to mainly employ a hydrogenotrophic lifestyle with many species being able to use formate (Imachi and Sakai 2015; Oren 2014). Some species from *Methanobacteriaceae* have been shown to use methanol with H_2_ (Fricke et al. 2006). Beside acetate, H_2_ is a major by-product of various fermentative processes and would be available to methanogens and other hydrogenotrophs in these sediments. Thus, the presence of all detected methanogens indicates a niche separation by availability of different substrates or/and fluctuations in acetate/H_2_ concentrations.

### Bacterial 16S rRNA gene distribution in the Bothnian Sea sediments

Similar to the archaeal communities, the bacterial 16S rRNA gene distribution between the coastal and central basin was significantly different (Figure 4). Both, N10 and NB8 were more similar to each other than NB8 and US5B (nMDS, Figure 5 (A)).

Particularly, the top sediment layer revealed substantial differences between the sites. At US5B, an apparent population of aerobic CH_4_ oxidizing bacteria (MOB) represented by *Methylococcaceae* was detected (6% of total bacterial reads). In contrast, this group was significantly lower in abundance at coastal sites with only few detected sequences. An enrichment of this group close to the sediment surface would point to a less efficient CH_4_ removal in deeper anoxic layers at US5B, particularly the SMTZ where most of the CH_4_ would be oxidized by ANME. CH_4_ that is not consumed in the SMTZ diffuses towards the sediment surface and fuels aerobic methanotrophic communities. The absence of surface sediment MOB communities at the coastal sites could be a result of either more efficient removal within the SMTZ (NB8) or a lower production by methanogens in deeper layers as seen at site N10.

The genus *Spirochaetales* was relatively more abundant at US5B, reaching a contribution of 12% in the deeper part of the profile. At both coastal sites, the abundances did not exceed 4%. In contrast, at both coastal sites *Anaerolineales* and *Bacteroidetes*_VadinHA17 were relatively more abundant than at US5B. All three groups, *Anaerolineales*, *Bacteroidetes*_VadinHA17 and *Spirochaetales* belong to an anaerobic core community involved in different fermentation pathways. The metabolic potential of *Anaerolineales*, reconstructed from several sequenced genomes and cultured representatives, points to a strictly anaerobic chemo-organotrophic lifestyle (Hug et al. 2013; Yamada et al. 2006). Members of the genus *Spirochaeta* have been shown previously to be an integral part of anoxic sediment communities (Breznak and Warnecke 2008; Shivani et al. 2015; and references therein). They are free-living, chemo-organotrophic facultative or obligate anaerobes capable of production of various fermentation products including acetate, ethanol, H_2_ and CO_2_ (Breznak and Warnecke 2008; Miyazaki et al. 2014). *Bacteroidetes*_VadinHA17 is an abundant member of sediment communities and has been discussed to be involved in degradation of organic matter (Bolhuis et al. 2014; Harrison et al. 2016). The observed differences in abundance of these three groups indicated that organic metabolite flows are different between coastal and central-basin sediments and could probably be explained by the quality of organic matter.

Another significant group of 16S rRNA gene sequences detected at all sites in different proportions was assigned to *Xanthomonadales*. Their sequences were already detected in high abundance at site US5B in our previous study (Rasigraf et al. 2017) and they are here shown to be ubiquitous in the Bothnian Sea sediments among analyzed depths. Sequences belonging to *Xanthomonadales* have previously been observed to be abundant in marine and brackish sediments (Dyksma et al. 2016; Mußmann et al. 2017). Based on the genomic information, they have been predicted to play an important in role S and N cycles by employing either chemolithoautotrophic or -heterotrophic lifestyle (Mußmann et al. 2017). Thus, *Xanthomonadales* may be a major contributor to dark CO_2_ fixation in marine sediments (Dyksma et al. 2016).

*Atribacteria* (formerly known as candidate divisions “OP9” and “JS1”) increased in relative abundance in the deeper part of the N10 sediment and reached up to 10% of the total bacterial 16S rRNA gene reads. At, NB8 and US5B, their abundance did not exceed 3% of total bacterial reads. *Atribacteria* have been shown previously to be abundant in anaerobic low energy environments (Newberry et al. 2004). Based on the available genome information, they appear to perform either primary fermentation, secondary fermentation or syntrophy for catabolism (Carr et al. 2015; Nobu et al. 2016). Their sequences have also been detected in Fe- and CH_4_-rich marine sediments of the Helgoland area in the North Sea (Oni et al. 2015a). There, their abundance strongly correlated with concentrations of dissolved Fe and CH_4_, and their possible involvement in Fe-dependent AOM together with members of *Methanosaetaceae* and the ANME-3 clade was suggested (Oni et al. 2015a). Other studies have also reported a regular occurrence of *Atribacteria* in sediments dominated by SO_4_^2−^-dependent AOM (Harrison et al. 2009). Carr et al. 2015 identified a strong correlation between dissolved CH_4_ profiles and abundance of *Atribacteria* in Arctic marine sediments. The observed correlation was suggested to be based on metabolic co-operation with methanogens which would scavenge fermentation products of *Atribacteria*, primarily acetate (Carr et al. 2015). The conditions at site N10 seem to favor the presence of *Atribacteria* in contrast to putative *Spirochaetales* fermenters at site US5B. *Desulfobacterales* were high in abundance at all sites. Members of *Desulfobacterales* include many characterized sulfate reducing bacteria (SRB) which use SO_4_^2−^ and other sulfur compounds as terminal electron acceptors and a variety of fermentation products as electron donors (Pfennig et al. 1981). At NB8 and US5B, a top to bottom gradient could be observed with highest abundances coinciding with the SMTZ. At N10, no apparent gradient could be observed and their 16S rRNA genes were distributed rather evenly over the whole sediment profile. At this site the SO_4_^2−^ penetration depth is also deeper than at NB8 and US5B (Figure 2). *Desulfobacterales* are often detected in marine sediments where SO_4_^2−^ and CH_4_ are present (Leloup et al. 2007; Ruff et al. 2015). Some members of the *Desulfobacterales* are frequently observed partners in ANME/SRB consortia, where they perform SO_4_^2−^ reduction and scavenge the reducing equivalents from ANME (Schreiber et al. 2010). Different ANME clades prefer certain SRB groups as partners, and it has been shown previously that ANME-2a are often detected together with SEEP-SRB1a – a clade belonging to *Desulfobacterales* (Schreiber et al. 2010). Our results are in line with previously published studies as the dominant ANME clade observed so far at all sites in the Bothnian Sea sediment belonged to ANME-2a. As expected, their highest abundance was observed in the zone where SO_4_^2−^ was detectable and where SO_4_^2−^ reduction was expected to occur. However, despite SO_4_^2−^ being under the detection limit (75 μM) below the SMTZ at NB8 and US5B, a zone where Fe-dependent CH_4_ oxidation was postulated to occur (Egger et al. 2015b), the presence of SRB was indicative of either a presence of a high flux of oxidized S-species or SRB performing other types of metabolisms (e.g. fermentation). Previous research has shown that SRB can switch from the respiratory metabolism to fermentation when suitable electron acceptors are not available (Plugge et al. 2011). In such situation, a cooperation with H_2_-scavenging methanogens is feasible (Plugge et al. 2011). Thus, a sudden introduction of SO_4_^2−^ could potentially activate their SO_4_^2−^ metabolism.

*Verrucomicrobiales* was abundant at both coastal sites but not in the central basin site US5B. Its distribution showed a strong gradient with highest numbers (up to 11% at NB8) near the sediment-water interface and a rapid decline within the sediment column. This change with depth points to an adaptation to high redox potential and possibly an aerobic/denitrifying lifestyle of dominant members making up the bulk of detected *Verrucomicrobiales* sequences. *Verrucomicrobia* have been previously shown to be abundant in marine water columns and sediments and to be mostly involved in polysaccharide degradation (Cardman et al. 2014; Martinez-Garcia et al. 2012). A similar distribution of *Verrucomicrobia* sequences was observed previously and was linked to degradation of fresh algal biomass in surface sediments of the North Sea (Oni et al. 2015b). Thus, based on previously available data, *Verrucomicrobiales* would belong to a community of primary degraders and possibly provide substrates for anaerobic fermentative communities.

Sequences belonging to *Flavobacteriales* were detected in high abundance at all sites. Highest abundances were observed within the SO_4_^2−^ penetration zone and SMTZ, similar to that of *Desulfobacterales*. At NB8, their abundance reached 10% of total bacterial reads within the SMTZ. At N10, their numbers slightly declined from top to bottom of the core with depth in the sediment, but did not exceed the maximum of 6%. Interestingly, at US5B, the *Flavobacteriales* sequence distribution initially followed that of both coastal sites with higher numbers at the top (6%) and declining towards 2% below the SMTZ, but the relative abundance started to increase again at the bottom of the sediment profile reaching up to 10% in the lowest analyzed depth of 52.5-55 cm. Interestingly, members of the *Flavobacteriales* were detected previously in an anaerobic methanotrophic enrichment originating from marine sediments (Jagersma et al. 2009). In active AOM cultures dominated by an ANME-2a archaeon, *Flavobacteriales* and *Desulfobacterales* together made up the bulk of the total bacterial sequences (Jagersma et al. 2009). Their metabolic role in that enrichment culture remained unclear, and possible involvement in S-compound transformations was discussed (Jagersma et al. 2009).

Another abundant group of bacteria detected at NB8 was assigned to the *Planctomycetales*. Their sequence abundance reached 6% and remained fairly constant throughout the sediment profile by minor variations between 3-6%. At N10, highest percentages of the total community were observed in the top 3 cm (6-7%), below which the population stayed at ca. 3% of the total bacterial reads. By further zooming in to a genus level, most reads were found to be assigned to the *Blastopirellula*, *Rhodopirellula*, *Bythopirellula* and Pir4_lineage. Some members of these lineages have been previously detected, described and isolated from Fe- and CH_4_-rich marine sediments (Storesund and Øvreås 2013; Winkelmann et al. 2010). They were shown to be involved in sugars and complex carbohydrate degradation, some were speculated to be involved in either Fe- or CH_4_ oxidation (Storesund and Øvreås 2013). Thus, the most abundant *Planctomycetales* residing in the Bothnian Sea sediment are most likely involved in the hydrolysis and degradation of complex organic matter.

### Metagenomic analysis of the Fe-rich methanic sediment at site NB8 in the Bothnian Sea

Metagenome assembly and binning of sediment samples at the coastal site NB8 resulted in a retrieval of 53 bacterial and 11 archaeal genomic bins with variable degree of completeness (Supplementary Table 1, only bins with >20% completeness, contamination level <10% and >0.1% proportion of total sequenced community were analyzed). In line with the abundance frequency in the 16S rRNA amplicon sequencing data, genome bins could be obtained for most abundant bacterial lineages including *Spirochaeta*, *Aminicenantes*, *Atribacteria*, *Chloroflexi, Actinobacteria, Bacteroidetes, Gemmatimonadales*, *Nitrospira*, *Planctomycetes, Parcubacteria, α-, β-, γ-, δ*-Proteobacteria and archaeal lineages including *Thaumarchaeota*, *Bathyarchaeota*, *Thorarchaeota*, *Methanomassiliicoccales*, *Methanosaeta, Methanosarcina,* ANME. We analyzed all bins and draft genomes for the presence of marker genes involved in fermentation, autotrophy/acetogenesis, methanogenesis/-trophy and respiratory N and S cycles (Figures 6 and 7, Supplementary Table 2).

**Figure 6:**
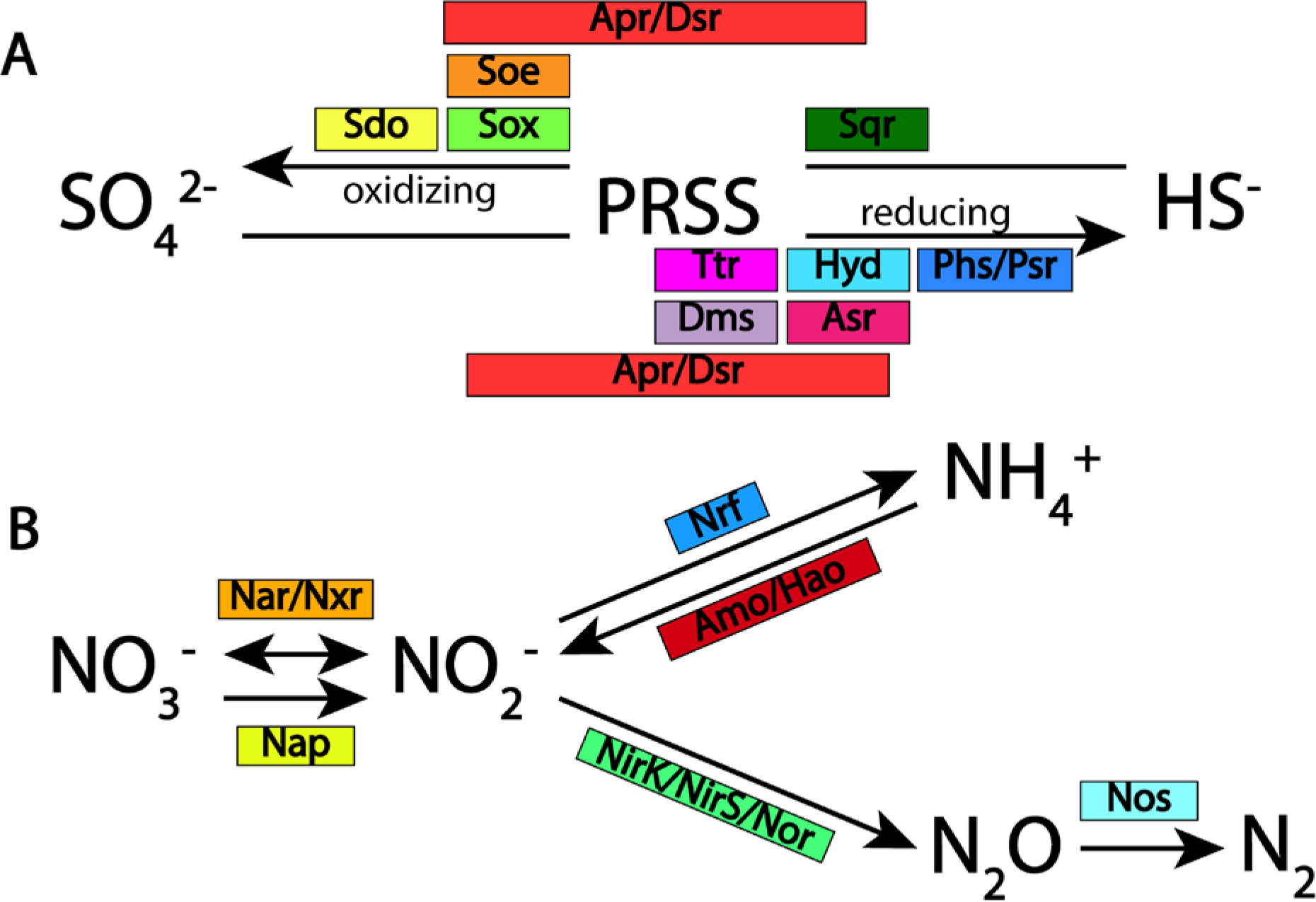
Metabolic potential for respiratory sulfur (a) and nitrogen (b) cycle reactions identified in metagenomic bins obtained from the iron-rich methanic sediment at site NB8 in the Bothnian Sea. Key enzymes catalyzing each process are shown. Abbreviations: PRSS, partially reduced sulfur species (include tetrathionate, thiosulfate, sulfite, polysulfide, elemental sulfur); Sdo, sulfur dioxygenase; Sox, sulfur-oxidizing multi-enzyme complex; Apr, adenylylsulfate (APS) reductase; Dsr, dissimilatory (bi)sulfite reductase; Sqr, sulfide:quinone oxidoreductase; Ttr, tetrathionate reductase; Asr, sulfite reductase; Hyd, sulfhydrogenase; Phs, thiosulfate reductase; Nar, nitrate reductase; Nxr, nitrate:nitrite oxidoreductase; Nap, periplasmic nitrate reductase; Nrf, nitrite reductase (NH_4_^+^ forming); Amo, ammonia monooxygenase; Hao, hydroxylamine oxidoreductase; Nos, nitrous oxide reductase; NirK, Cu-containing nitrite reductase (NO forming); NirS, Fe-containing nitrite reductase (NO forming); Nor, nitric oxide reductase.

**Figure 7:**
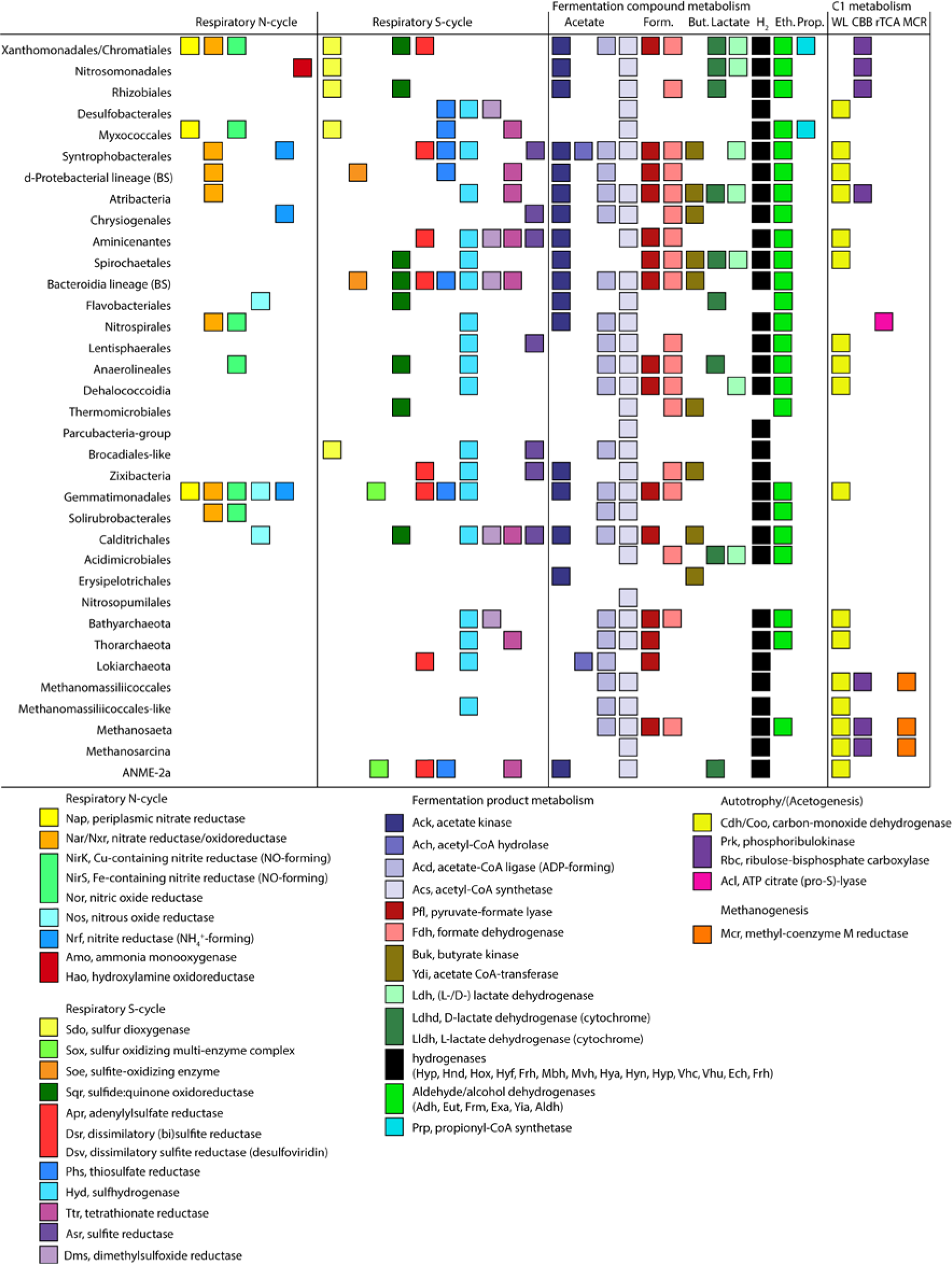
Overview for presence of functional biomarkers in genome bins obtained from the Fe- rich methanic sediment layer at coastal site NB8 in the Bothnian Sea. Analysis was performed for key genes encoding enzymes involved in processes of respiratory nitrogen (N) and sulfur (S) cycles, fermentation product metabolism, autotrophy/acetogenesis and methanogenesis. Abbreviations: Form., formate; But., butyrate; Eth., ethanol; Prop., propionate; WL, Wood-Ljungdahl pathway; CBB, Calvin-Benson-Bessham cycle; rTCA, reverse tricarboxylic acid cycle.

### Respiratory sulfur (S) cycle metabolism

In anaerobic sediments, Fe and manganese (Mn) oxides can undergo abiotic reactions with H_2_S and lead to its oxidation to either partially reduced sulfur species (PRSS, comprising thiosulfate, polysulfide, tetrathionate, sulfite and elemental sulfur) or completely to SO_4_^2−^ (Zopfi et al. 2004). Anoxic Bothnian Sea sediments below the SMTZ have been shown to contain high concentrations of Fe oxides which consist for >50% of ferric (oxy)hydroxide (Egger et al. 2015a; Lenstra et al. 2018; Slomp et al. 2013). Thus, any free H_2_S is likely to react fast either with the Fe oxides or precipitate as FeS with Fe^2+^. Partial oxidation of free H_2_S with Fe oxides has been suggested to lead to formation of SO_4_^2−^ and thiosulfate (or other PRSS) which would act as an electron acceptor source for organisms which would reduce the PRSS and SO_4_^2−^ with donors such as acetate or H_2_ back to H_2_S (Zopfi et al. 2004). This cycle has been described as cryptic S cycle in marine anoxic sediments (Brunner et al. 2016; Holmkvist et al. 2011). We analyzed the presence of functional gene biomarkers involved in reductive processes such as thiosulfate/polysulfide reductase (Phs/Psr), sulfhydrogenase (Hyd), tetrathionate reductase (Ttr), sulfite reductase (Asr), dimethyl sulphoxide (DMSO) reductase (Dms), adenylyl-sulfate reductase (Apr)/dissimilatory (bi)sulfite reductase (Dsr). The latter, Apr/Dsr complex, was also shown to catalyze the reverse reaction of H_2_S oxidation to SO_4_^2−^ in some *Proteobacteria* and *Chlorobi* (Ghosh and Dam 2009; Müller et al. 2015).

In general, genes potentially involved in PRSS transformations were detected in most of the retrieved bacterial MAGs indicating a potential for an active S cycle in the methanic zone below the SMTZ. Among archaeal MAGs, the most widespread PRSS metabolism biomarkers encoded sulfhydrogenase-like proteins. However, genes encoding all four subunits (HydABDG) were only detected in bins assigned to *Thorarchaeota*, corroborating recent findings about these recently characterized organisms (Seitz et al. 2016). Other bacterial and archaeal MAGs mostly only encoded one or two of the four Hyd-like comprising subunits.

Biomarkers for dissimilatory SO_4_^2−^ reduction to H_2_S (Apr/Dsr) were detected in six bacterial MAGs. Apr without the presence of Dsr was found in *Bacteroidales* and *Xanthomonadales*/*Chromatiales*, while Dsr without Apr was detected one *Aminicenantes* MAG. Both biomarkers were detected in *Syntrophobacterales* and *Gemmatimonadales*. Many members of the *Syntrophobacterales* order are SO_4_^2−^ reducers frequently detected in anaerobic SO_4_^2−^ containing sediments (Plugge et al. 2011). The finding of one *Gemmatimonadales* genomes also containing both Apr and Dsr encoding genes was more surprising. Recently, similar observations were reported for *Gemmatimonadales* MAGs obtained from estuarine sediments (Baker et al. 2015), however no SO_4_^2−^ reducers have been described so far from this group. Thus, these previously unknown potential SRB might be widespread in estuarine, marine and brackish sediments and their role in SO_4_^2−^ reduction might have been overlooked in the past.

The detected Apr in one the *Xanthomonadales*/*Chromatiales* MAGs indicated potential involvement in sulfite oxidation. Members of *Chromatiales* and particularly *Ectothiorhodospiraceae* family, have been frequently detected in marine anoxic sediments and were shown to employ a chemolithoautrophic lifestyle of either Fe- or reduced S compound oxidation (Dyksma et al. 2016; Hallberg et al. 2011). The detection of a phosphoribulokinase (Prk) in one of the MAGs classified into this group further pointed to some of them likely being autotrophs.

Interestingly, homologues of a desulfoviridin-type dissimilatory sulfite reductase were detected in MAGs classified as ANME-2a and *Lokiarchaeota*. This type of a sulfite reductase is involved in an energy-yielding reduction of sulfite to H_2_S. This finding is particularly interesting in the view of anaerobic CH_4_ oxidation potential in these sediments. ANME are typically associated with SRB in order to perform AOM, in which the bacterial partner would perform the reduction of SO_4_^2−^ or PRSS to H_2_S (Knittel and Boetius 2009). However, this finding indicates the potential of Bothnian Sea ANME to perform the reduction of sulfite intrinsically. The ability of some ANME to reduce sulfur species has been observed earlier (Milucka et al. 2012).

Several core community members of putative fermenters including *Anaerolineales*, *Bacteroidales*, *Atribacteria* and *Aminicenantes* also encoded gene homologues for enzymes involved in PRSS transformations. This indicated flexible metabolic strategies switching between PRSS respiration and fermentations depending on environmental conditions.

Also DMSO reductase-like encoding genes were detected in several retrieved MAGs. DMSO is a common metabolite in marine and brackish ecosystems were it is produced by microalgae, phytoplankton and angiosperms as osmoprotectant (López and Duarte 2004). DMSO can then be used as electron acceptor under anaerobic conditions which results in the production of dimethyl sulfide. Among others, one MAG assigned to *Aminicenantes*, which is based on metagenome data the most abundant bacterial group in the investigated sediment layer (Supplementary Table 1), contained Dms-like encoding genes. Thus, potential for reductive respiratory S cycle metabolisms seems widespread in core community bacterial taxa.

In the presence of energetically more favorable electron acceptors like NO_3_^−^ or Mn^4+^, reduced sulfur compounds can be completely oxidized to SO_4_^2−^ (Zopfi et al. 2004). This process would represent a source of SO_4_^2−^ and thus electron acceptor for SRB. The analyzed biomarkers for PRSS and H_2_S oxidation included sulfur dioxygenase (Sdo), sulfur oxidizing multi-enzyme system (Sox), sulfide:quinone oxidoreductase (Sqr) and sulfite oxidase (Soe). In general, homologues of genes encoding enzyme subunits involved in oxidative processes in the S cycle were less widespread than those involved in PRSS reduction. This redundancy in the potential for oxidative processes could be further explained with the lack or shortage of electron acceptors in this sediment layer. Thus, the residing microbial community would over time lose the ability for PRSS oxidation.

The results of respiratory S cycle analysis indicated the potential for PRSS reductive processes in many analyzed MAGs while that for oxidative ones was scarcer. These findings point to a possibility of a shorter operational S cycle where the abiotic oxidation of H_2_S by the reactive Fe would create a pool of PRSS which would be reduced back to H_2_S.

### Respiratory nitrogen (N) cycle

N cycle activities have been recently investigated in Öre Estuary sediments (Hellemann et al. 2017). Nitrification-denitrification seemed to be the dominant sink for reactive N in the ecosystem while anaerobic ammonium oxidation (anammox) was not detectable (Hellemann et al. 2017). Several genomic bins obtained within this study contained gene homologues encoding various enzymes catalyzing respiratory N cycle processes.

For the initial step of NO_3_^−^ reduction, two types of NO_3_^−^ reduction systems were analyzed: a periplasmic (Nap) and membrane-bound nitrate reductase (Nar). Nar resembles high similarity to nitrite oxidoreductase (Nxr) which catalyzes oxidation of nitrite (NO_2_^−^) to NO_3_^−^. Both Nap and Nar exhibited similar abundance among analyzed bacterial taxa. Only two groups, *Xanthomonadales*/*Chromatiales* and *Gemmatimonadales* contained both Nap and Nar. In addition, some *Gemmatimonadales* MAGs also contained genes encoding for enzymes catalyzing reduction of NO_2_^−^ to nitric oxide (NO) (NirK) and reduction of nitrous oxide (N_2_O) to N_2_ (Nos). Next to a wide spectrum of genes involved in fermentation product metabolism, SO_4_^2−^ and PRSS reduction, the order *Gemmatimonadales* appeared to possess high metabolic diversity.

Nxr was only detected in one MAG classified as *Nitrospirales*, an order which includes many characterized NO_2_^−^ oxidizers widespread in natural ecosystems (Lücker et al. 2010). Additionally, it also contained a Cu-dependent nitrite reductase (NirK) encoding gene. Possibly, the detected *Nitrospira* organisms could switch between NO_2_^−^ oxidation and denitrification depending on redox potential and substrate availability.

*Syntrophobacterales* encoded a Nar and cytochrome c nitrite reductase (Nrf), besides the potential for SO_4_^2−^ reduction. Previous studies with characterized SRB have shown the preferred use of NO_3_^−^ as an electron acceptor when available (Krekeler and Cypionka 1995). The capability for DNRA seems to be a common trait among SRB and indicates a flexible metabolism depending on the availability of electron acceptors and a coupling of dissimilatory N and S cycles. Another recent study showed that SRB *Desulfurivibrio alkaliphilus* can employ a chemolithotrophic metabolism by NO_3_^−^ reduction to NH_4_^+^ and oxidation of H_2_S to thiosulfate or elemental sulfur (Thorup et al. 2017). Research on partnerships of ANME with different types of SRB has indicated that availability of NO_3_^−^ may play an important role in the establishment of certain types of ANME/SRB symbioses (Green-Saxena et al. 2014). NO_3_^−^ was shown to be used as an N source, however its incorporation into biomass was secondary to NH_4_^+^ and the authors could not exclude DNRA as a possible mechanism (Green-Saxena et al. 2014).

Functionally, dissimilatory NO_3_^−^ reduction can be decoupled from further steps of NO_2_^−^ reduction and an organism can excrete NO_2_^−^ which can be further used as an electron acceptor by other community members. The fate of NO_2_^−^ then differs depending on genomic potential. It can be either reduced to NO and then to N_2_O or reduced in one step to NH_4_^+^ by Nrf. NO is a toxic and very reactive metabolite which is usually processed by the cell immediately. Thus, we analyzed the presence of NO-forming nitrite reductases (NirK/NirS) in combination with NO reductases (Nor) as one module for denitrification to N_2_O. The product of this process is N_2_O which again can be either excreted into the environment or further reduced to N_2_. N_2_O reduction can be performed by a different functional group of denitrifiers. Both denitrification modules, to N_2_O and to N_2_, were spread among the retrieved MAGs indicating functional redundancy and truncation in metabolic denitrification potential. However, activity measurements performed by Hellemann et al. 2017 revealed that the contribution of N_2_O production to total denitrification in Öre Estuary sediments was below 1% and was thus negligible. The ratio of N_2_O/N_2_ as end product of denitrification is influenced by several factors including organic carbon and NO_3_^−^ loads (Weier et al. 1993). The Öre Estuary is an oligotrophic system which is limited in easy accessible electron donors (Hellemann et al. 2017). However, seasonal changes in the input of organic matter and N availability might shift the ratio of N_2_O to N_2_ production in the system.

The potential for DNRA was only present in *Syntrophobacterales, Gemmatimonadales* and *Chrysiogenales* MAGs. They also possessed marker genes for PRSS transformations, thus DNRA could be driven by electrons derived from the oxidation of H_2_S or alternatively fermentation products. The activity of either DNRA or denitrification for NO_2_^−^ reduction would depend on the quality and availability of electron donors in the estuary sediment system.

Nitrification potential was assessed by the presence of Amo/Hao encoding genes. Amo/Hao catalyzes the oxidation of ammonia to NO_2_^−^, which can then be used by NO_2_^−^ oxidizers such as *Nitrospira* for further oxidation to NO_3_^−^ by an Nxr. Both processes require a potent electron acceptor such as oxygen (O_2_). Two MAGs assigned to putative ammonia oxidizers could be retrieved from the analyzed sediment. One was assigned to *Nitrosomonadales*, a bacterial order which includes many characterized ammonia oxidizers, and another to archaeal *Thaumarchaeota*.

However, only the *Nitrosomonadales* MAG contained both Hao and Amo, the thaumarchaeal MAG lacked the genes encoding for Hao/Amo. However, since the latter was only 46% complete, it is likely that ammonia oxidation pathway encoding genes were not binned into the MAG. Both ammonia oxidizing bacteria (AOB) and archaea (AOA) might be participating in the oxidation of NH_4_^+^ to NO_2_^−^ in the coastal Bothnian Sea sediment. It is not clear, however, how O_2_ would be available for their metabolism in the analyzed depth since air is unlikely to penetrate to the methanic zone below the SMTZ. It has been shown previously that O_2_ penetration depths are restricted to the upper centimeter in the Öre Estuary sediments (Hellemann et al. 2017). The presence of aerobic nitrifiers in anoxic environments has been frequently observed in the past and an alternative anaerobic metabolism was discussed as a possible lifestyle strategy (Abeliovich and Vonshak 1992; Schmidt et al. 2002; Weber et al. 2001). Some studies from the 1990s and more recent ones have hypothesized the possibility of nitrification coupled to metal oxide reduction involving, for example, Fe and Mn oxides (Hulth et al. 1999; Luther et al. 1997; Mogollón et al. 2016; Thamdrup and Dalsgaard 2000). An alternative explanation would be a dormant nitrifier community which was preserved at this depth due to fast sedimentation and slow degradation rates.

Our results show that even in the complete absence of O_2_ and despite the activity of alternative anaerobic processes, the analyzed sediment still contains a genetic potential for O_2_-dependent nitrification.

### Fermentative metabolism

Fermentative processes are of central importance in sediment ecosystems since anaerobic degradation of deposited organic matter yields a variety of short chain fatty− and carboxylic acids and H_2_ which can be further used in respiratory processes for the reduction of oxidized N-, S− and Fe species, methanogenesis and homoacetogenesis (Finke et al. 2007). The production and consumption of those organic and inorganic (H_2_) intermediates depends on factors such as sediment pH, temperature, quality of the deposited organic matter and availability of inorganic electron acceptors. These factors are expected to vary depending on seasonality, external input variability, bioturbation and sedimentation rates. Thus, the presence of gene biomarkers only represents the potential of the system for the analyzed processes and not the actual metabolite flows.

In marine and brackish sediments where the pH is usually between 7 and 7.5, major fermentation products comprise acetate, formate, ethanol, propionate, butyrate, lactate and H_2_. Acetate is the central metabolite in marine sediments (Shaw and McIntosh 1990). We analyzed the presence of four functional biomarkers involved in acetate metabolism: (ADP-forming) acetate-CoA ligase (Acd), acetate kinase (Ack), acetyl-CoA hydrolase (Ach) and acetyl-CoA synthetase (Acs). Those enzymes can catalyze reactions in both directions which will depend on environmental conditions and the employed metabolism by the organism.

In general, most MAGs contained the genes encoding either one of several of the abovementioned enzymes. Acd and Acs were the most widespread acetate metabolism biomarkers among both bacteria and archaea including *Thorarchaeota*, *Bathyarchaeota* and *Lokiarchaeota*. Thus, those archaea could contribute to fermentative acetate production or assimilation in methanic sediments below SMTZ.

Formate metabolism was assessed by the presence of pyruvate-formate lyase (Pfl) which catalyzes formate formation from pyruvate, and formate dehydrogenase (Fdh/Fdo) for formate oxidation. Both genes were widely distributed among bacterial and archaeal bins. Many putative fermenters including *Spirochaetales*, *Bacteroidales*, *Aminicenantes* and *Atribacteria* contained both. Among archaeal MAGs, the potential for acetate and formate turnover was widespread corroborating previous results for *Bathyarchaeota* and *Thorarchaeota* being potentially involved in fermentative production of acetate and formate (Lazar et al. 2016; Seitz et al. 2016). The lowest distribution was observed for propionate turnover encoding genes (Prp).

The ability for ethanol metabolism which was assessed by the presence of aldehyde and alcohol dehydrogenases (Aldh/Adh/Exa/Frm/Yia/Eut) appeared to be one of the most widespread traits among the retrieved bacterial MAGs. This was an indication for ethanol being next to acetate and formate an important metabolite in the analyzed sediment system. Among archaea, however, it was only detected in *Bathyarchaeota*, *Thaumarchaeota* and *Methanosaeta*, indicating their potential involvement in ethanol production/uptake.

Lactate metabolism was assessed by the presence of cytochrome-(Lldh/Ldhd) and NAD(P)-dependent (Ldh) lactate dehydrogenase encoding genes. Several bacterial MAGs possessed lactate utilization biomarkers. In archaea only the highly incomplete MAG assigned to ANME contained an Lldh-like gene. The rest of archaeal bins did not seem to possess capacity for lactate metabolism.

The ability for H_2_ metabolism was assessed by the presence of genes encoding for subunits of various types of hydrogenases. Genes encoding for the following hydrogenase complexes were detected in the analyzed MAGs: bi-directional NAD(H)-dependent [NiFe] hydrogenase (Hox), periplasmic [NiFeSe] hydrogenase (Hya), [Fe] hydrogenase (Hyd), [NiFe] hydrogenases which couple H_2_ production to formate or CO oxidation, ferredoxin-dependent bi-directional [NiFe] hydrogenase Ech, [FeFe] hydrogenases mostly involved in H_2_ production but also oxidation in SRB particularly and F420-non-reducing hydrogenases. Overall, genes encoding hydrogenase subunits or hydrogenase maturation pathways were detected in most of the analyzed bacterial and archaeal MAGs indicating the central importance of H_2_ in the analyzed sediment ecosystem.

Putative bacterial fermenters including *Spirochaetales*, *Bacteroidales*, *Anaerolineales*, *Aminicenantes* and SRB contained genes encoding several hydrogenase systems indicating an adaptation to ambient fluctuations in metabolite concentrations. One of the *Gemmatimonadales* MAGs which encoded the whole SO_4_^2−^ reduction pathway also contained genes encoding for two types of hydrogenases. Thus, this potential SRB could be utilizing H_2_ as an electron donor for SO_4_^2−^ reduction. Similarly, also *Syntrophobacterales* which encoded the full SO_4_^2−^ reduction pathway revealed a wide H_2_ utilization potential via different hydrogenases. In general, all genomes with a Dsr encoded for one or several hydrogenases. Interestingly, also most archaeal MAGs contained genes encoding several hydrogenases indicating their important role in H_2_ metabolism in Bothnian Sea sediments.

### CO_2_ fixation and acetogenesis

Next to the organic matter input which is eventually turned over into CO_2_, energy and new biomass by heterotrophic organisms, CO_2_ fixation by autotrophs represents another organic carbon input route into the sediment ecosystem. In deep anoxic sediments, autotrophs usually gain energy from the oxidation of inorganic electron donors such as H_2_S, PRSS, H_2_ and reduced metals. In natural systems, CO_2_ can be fixed via several pathways (Berg 2011). We analyzed the presence of gene biomarkers for the Wood-Ljungdahl pathway (WL) (carbon monoxide dehydrogenase, Cdh/Coo), Calvin-Benson-Bassham cycle (CBB) (phosphoribulokinase (Prk) and ribulose-1,5-bisphosphate carboxylase (Cbb)) and reductive citric acid cycle (rTCA) (ATP citrate lyase (Acl)).

Here, the WL pathway is not only indicative of autotrophy, but can also be used for acetate production by acetogens or assimilation of acetate, CO, or methylamines (Berg 2011). Methanogenic archaea use the WL pathway for both autotrophic CO_2_ fixation and methanogenesis (Berg 2011). The Cdh/Coo biomarker was widespread among both bacterial and archaeal MAGs. As expected, the putative *Syntrophobacterales* SRB and other *δ*-proteobacterial MAGs contained Cdh/Coo biomarkers. Previous research showed that SRB utilize the WL pathway in both reductive and oxidative directions (Schauder et al. 1988). One of the bins classified as *Lentisphaerales* contained Cdh/Coo biomarkers. So far, no reports on the presence of the WL pathway in these organisms are available. Thus, they might represent novel acetate producers/scavengers in methanic sediments.

Also putative fermenters including *Spirochaetales*, *Anaerolineales*, *Aminicenantes* and *Atribacteria* contained Cdh/Coo-encoding biomarkers. In fermenters, the WL pathway was discussed to function as an electron sink by reduction of CO_2_ to acetate (Berg 2011). Thus, those organism groups might be producing acetate via this route during fermentation. The potential for the WL pathway has also been detected previously in the MAG of a putatively fermentative *Chloroflexi* bacterium RBG-2 (Hug et al. 2013) and sediment-derived genomic bins assigned to *Anaerolineales* (Fullerton and Moyer 2016). It was discussed to be operating either under heterotrophic conditions to reduce the intracellular CO_2_ by simultaneous oxidation of reduced ferredoxin and NADH, or under autotrophic conditions for CO_2_ fixation (Hug et al. 2013; Ragsdale and Pierce 2008).

In many of the same MAGs which contained Cdh/Coo biomarkers we detected genes encoding for pyruvate-ferredoxin oxidoreductase (Pfor) which might point to a link between autotrophic CO_2_ fixation via WL pathway and TCA cycle via acetyl-CoA (Furdui and Ragsdale 2000). Cdh/Coo biomarkers were widespread among archaeal MAGs including methanogens, *Bathyarchaeota* and *Thaumarchaeota*. As discussed previously, methanogens use an archaeal variant of the WL pathway which is employed in hydrogenotrophic and acetotrophic methanogenesis (Berg 2011; Borrel et al. 2016). Interestingly, we also detected Cdh/Coo in one retrieved *Methanomassiliicoccales* MAG. To date, *Methanomassiliicoccales* methanogens, which have only been described recently, were collectively implicated in lacking the WL pathway and thus being restricted to H_2_-dependent methylotrophic methanogenesis (Borrel et al. 2016). By further GenBank protein database search we found that at least two other *Methanomassiliicoccales* genomes (RumEn M1 and RumEn M2) encode Cdh/Coo biomarkers (accession nr. KQM11260 and KQM09953). Thus, some *Methanomassiliicoccales* genomes encode parts of the WL pathway. The detection of Cdh/Coo and other enzymes of the archaeal-type methylotrophic branch of WL pathway in one of the *Bathyarchaeota* MAGs obtained in this study further corroborated previous findings for this group of archaea (He et al. 2016; Lazar et al. 2016). *Bathyarchaeota* have been discussed to employ WL pathway for acetogenesis (He et al. 2016; Lazar et al. 2016). Similarly, Cdh/Coo and other genes encoding for the archaeal variant of the WL pathway were present in *Thorarchaeota* MAGs which hints to their involvement in acetogenesis in the analyzed sediment system.

Next to the WL pathway which is mainly found in anaerobic organisms operating close to the thermodynamic limit, CBB cycle is employed by a variety of chemolithoautotrophic organisms and can operate under higher redox potentials (Berg 2011). We analyzed the presence of two biomarkers which are unique to the CBB cycle: Prk and Cbb. Among bacterial MAGs, Cbb in combination with Prk was only detected in *Xanthomonadales/Chromatiales* and *Nitrosomonadales*. The ability for autotrophic CO_2_ fixation via the CBB cycle has been recently reported to be widespread among *γ*-proteobacterial lineages *Woeseiaceae*/JTB255 which belong to the core community in diverse marine sediments (Mußmann et al. 2017) and to which Bothnian Sea *Xanthomonadales*/*Chromatiales* were closely related. Several genomes have been shown to encode biomarkers of CBB cycle, truncated denitrification pathway to N_2_O and PRSS oxidation to SO_4_^2−^ (Dyksma et al. 2016; Mußmann et al. 2017). *Xanthomonadales*/*Chromatiales* MAGs obtained from the Bothnian Sea sediment contained biomarkers for PRSS transformations, denitrification and CBB cycle. Thus, these ubiquitous *γ*-Proteobacteria could be involved in chemolithoautotrophic PRSS oxidation coupled to denitrification to N_2_O in the coastal methanic sediments of the Bothnian Sea. The detection of CBB biomarkers in the obtained *Nitrosomonadales* MAG was in accordance with characterized chemolithoautotrophic metabolism of this organism group (Utåker et al. 2002). However, as *Nitrosomonadales* might be involved in an alternative anaerobic metabolism in the analyzed sediment, the functionality of their CBB pathway remains unknown. This could be elucidated by future transcriptomic studies on this ecosystem.

Among archaea, Cbb encoding genes were detected in methanogens. Various methanogens have been shown previously to possess Cbb biomarkers, however their functionality remained debated. Recently, a functional pathway involving Cbb and Prk, similar to CBB cycle in autotrophic organisms, was proposed for methanogens (Kono et al. 2017). However, the ability for autotrophy based on this pathway among methanogens remains unclear (Kono et al. 2017).

rTCA cycle biomarker Acl was detected in one of the *Nitrospirales* MAGs which was closely related to a *Nitrospira* bacterium. NO_2_^−^-oxidizing *Nitrospira* have been reported previously to employ rTCA cycle for CO_2_ fixation (Lücker et al. 2010). Thus, this autotrophic pathway seems to be restricted to only one dominant group of bacteria residing in the analyzed sediment.

### Methanogenesis-/trophy

Three of the obtained archaeal MAGs could be classified as methanogens: *Methanosaeta*, *Methanosarcina* and *Methanomassiliicoccales*. Methyl-coenzyme M reductase (Mcr) was either partially or fully encoded in all three genomes. Members from those three groups have been described previously and represent different functional groups within methanogens. *Methanosaeta* from the family *Methanosaetaceae* is an obligate acetotroph which was reported to possess a high affinity to acetate and thrive under low ambient acetate concentrations (Jetten et al. 1992). In contrast, *Methanosarcina* methanogens possess a wide substrate spectrum but low affinity to acetate (Jetten et al. 1992). Despite high incompleteness of the obtained *Methanosarcina* genome, we identified genes encoding several complexes involved in methylotrophic metabolism. Both, *Methanosarcina* and *Methanosaeta* are abundant core community members in anaerobic methanic sediments (Carr et al. 2017; Webster et al. 2015). In contrast, the distribution of *Methanomassiliicoccales* methanogens in natural methanic sediments is underexplored. Originally, all described *Methanomassiliicoccales* were isolated or enriched from intestinal tracts of animals (Dridi et al. 2012). Since then, biomarkers of *Methanomassiliicoccales* have been detected in various sediment ecosystems and their distribution was investigated in more detail recently (Becker et al. 2016; Speth and Orphan 2018). All physiological and genomic information available so far point to a strictly H_2_-dependent methylotrophic methanogenesis by a complete lack of the archaeal type WL pathway (Borrel et al. 2016). The genome we obtained from the methanic sediment in this study contains Cdh/Coo biomarkers which encode one of the key genes in WL pathway. Thus, its role in metabolism remains unknown or it might represent a remnant of the WL pathway which has been lost during *Methanomassiliicoccales* evolution. Interestingly, one archaeal genome (metabat2.27) obtained from the Bothnian Sea sediment was closely related to available *Methanomassiliicoccales* genomes obtained from GenBank but lacked Mcr and other essential methanogenic biomarkers. It is unclear whether this organism is a methanogen and further analysis of this potentially novel organism group is needed.

16S rRNA gene amplicon sequencing data revealed that methanotrophic ANME-2 archaea were among the most abundant groups of archaea in the analyzed depth intervals. However, only one highly incomplete (22.6%) MAG classified as ANME-2a could be retrieved from the analyzed sediment. The MAG did not contain Mcr biomarkers so we analyzed the metagenome for its total *mcrA* inventory by blastx analysis (Figure 8). The result revealed ANME-like and *Methanosarcina*-like *mcrA* gene reads to make up the majority of the total *mcrA* pool.

**Figure 8:**
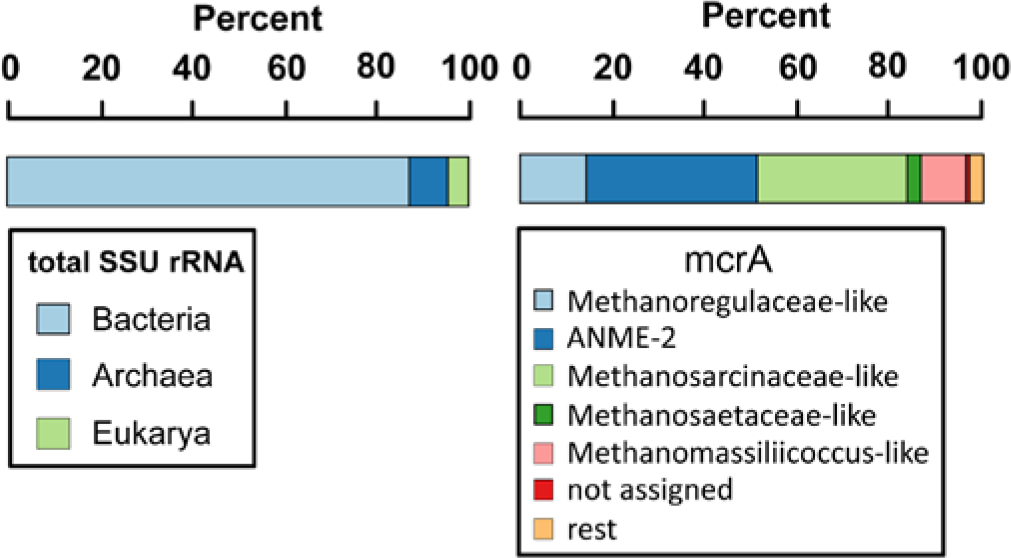
Domain distribution (left) and estimation of community diversity (right) in the Fe-rich methanic zone of site NB8 in the Bothnian Sea. Complete 16S rRNA and *mcrA* gene sequences were obtained from the assembled metagenome.

Our results show that methanogenesis in the coastal Bothnian Sea sediments would be mostly based on methylated compounds, acetate and less on CO_2_/H_2_. Methylated compounds are common substrates in marine and brackish sediments as they are degradation products of osmoregulators such as trimethylamine oxide and quaternary amines (Sørensen and Glob 1987). These compounds are the so-called non-competitive substrates for methanogens, as SO_4_^2−^ reducers do not use them for SO_4_^2−^ reduction (King 1984). In contrast, acetate and H_2_ are common electron donors for both SO_4_^2−^ reducers and methanogens so that methanogens are often outcompeted due to unfavorable substrate affinities (Oremland and Polcin 1982).

The potentially methanotrophic ANME-2a archaea could undergo interactions with electron-scavenging bacteria which would respire PRSS or Fe oxides, since many bacteria possess sulfur-based metabolisms. Or they would alternatively reduce SO_4_^2−^ themselves as indicated by the presence of a desulfoviridin-type sulfite reductase.

## Conclusions

The obtained genomes and total functional gene analysis of dominant organisms from the coastal methanic sediments in the Bothnian Sea indicated wide genetic potential for respiratory S cycle via PRSS transformations and diverse fermentation metabolisms with acetate, alcohols and H_2_ being potentially the major metabolites in the system. The potential for the respiratory N cycle was dominated by denitrification over DNRA and methanogenesis would be mostly based on methylated compounds and acetate. The archaeal population was dominated by putative anaerobic methanotrophs from the ANME-2a clade. Other abundant archaea which dominated the Bothnian Sea sediments included *Bathyarchaeota*, *Thorarchaeota* and *Lokiarchaeota*. Their genetic potential indicated fermentations and PRSS respiration as possible lifestyle strategies. All these processes will mainly depend on the quality and input amount of complex organic matter which will form the basis for the food chain in anaerobic sediments. This hypothesis is feasible since we observed significant differences in both archaeal and bacterial populations with distance from the shoreline in the Bothnian Sea despite the similar overall concentrations of major electron acceptors (e.g. Fe, SO_4_^2−^). With the PRSS, Fe oxides are thought to play a critical role in the observed redox transformations (Figure 9). For future studies enrichments and physiological characterizations of dominant communities would improve our understanding of their physiology, in situ biological function and biogeochemical impact.

**Figure 9:**
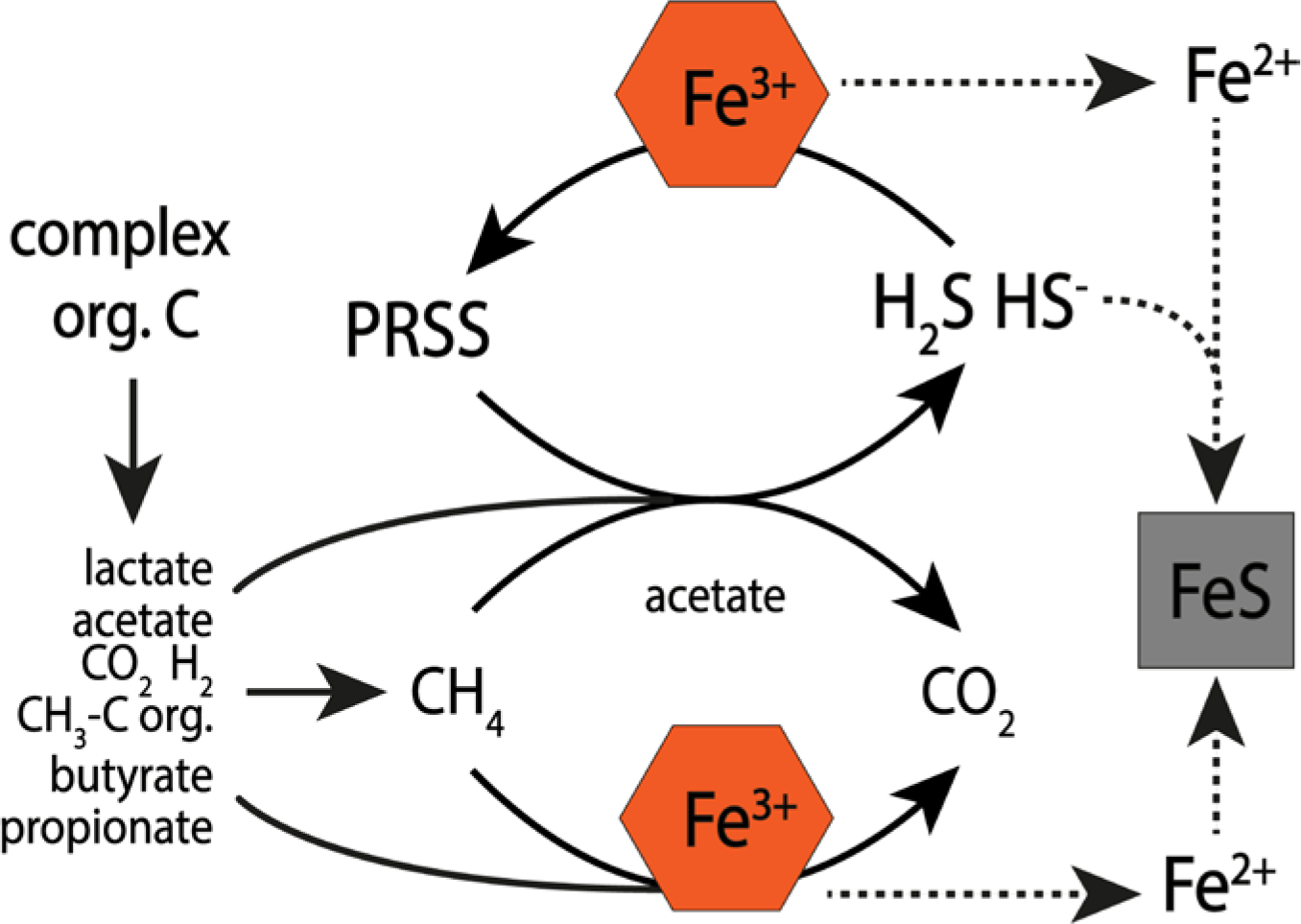
Simplified overview of predicted processes involving Fe and sulfur to take place in the analyzed coastal anaerobic sediments in the Bothnian Sea at site NB8. Ferrihydrite would represent the major electron acceptor in the analyzed sediments and be involved in the biotic and abiotic oxidation of sulfide and fermentation products. Anaerobic oxidation of methane could potentially be fueled by the reduction of PRSS formed by ferrihydrite or ferrihydrite directly. Acetate would be one of the key metabolites formed during the primary and secondary fermentations. The reduced Fe would react with the free sulfide and form insoluble iron sulfides. Abbreviations: PRSS, partially reduced sulfur species

## Acknowledgements

We would like to thank Theo van Alen and Geert Cremers for sequencing the metagenomes. The captain, crew and scientific participants aboard R/V *Lotty* are thanked for their assistance during sampling in August 2015. This work was carried out on the Dutch national e-infrastructure with the support of SURF Cooperative. LABGeM (Genoscope, Institute of Genomics, CEA Sciences) and the National infrastructure “France Génomique” are acknowledged for support within the MicroScope annotation platform. O.R., M.S.M.J. and C.P.S. were supported by NESSC (grant number 024002001), J.F. and M.S.M.J were supported by the SIAM Gravitation Grant on Anaerobic Microbiology (Netherlands Organization for Scientific Research, SIAM 024 002 002) and ERC AG (nr. 339880), C.P.S., W.L. and N.A.G.M.v.H. were supported by NWO (grant number 865.13.005) and the European Union and FORMAS through BONUS COCOA (grant number 2112932-1).

